# *Syntrophus* Conductive Pili Demonstrate that Common Hydrogen-Donating Syntrophs can have a Direct Electron Transfer Option

**DOI:** 10.1101/479683

**Authors:** David J.F. Walker, Kelly P. Nevin, Dawn E. Holmes, Amelia-Elena Rotaru, Joy E. Ward, Trevor L. Woodard, Jiaxin Zhu, Toshiyuki Ueki, Stephen S. Nonnenmann, Michael J. McInerney, Derek R. Lovley

## Abstract

Syntrophic interspecies electron exchange is essential for the stable functioning of diverse anaerobic microbial communities. Hydrogen/formate interspecies electron transfer (HFIT), in which H_2_ and/or formate function as diffusible electron carriers, has been considered to be the primary mechanism for electron sharing because most common syntrophs were thought to lack biochemical components, such as electrically conductive pili (e-pili), necessary for direct interspecies electron transfer (DIET). Here we report that *Syntrophus aciditrophicus*, one of the most intensively studied microbial models for HFIT, produces e-pili and can grow via DIET. Pilin genes likely to yield e-pili were found in other genera of hydrogen/formate-producing syntrophs. The finding that DIET is a likely option for diverse syntrophs that are abundant in many anaerobic environments necessitates a reexamination of the paradigm that HFIT is the predominant mechanism for syntrophic electron exchange within anaerobic microbial communities of biogeochemical and practical significance.

## Introduction

H_2_/formate interspecies electron transfer (HFIT) and direct interspecies electron transfer (DIET) are mechanisms for the electron-donating partner in syntrophic consortia to dispose of electrons released from the oxidation of key intermediates (organic acids, alcohols, aromatics) during the anaerobic degradation of complex organic matter ^1–5^. The relative proportion of electron flux through DIET or HFIT can influence the speed of interspecies electron transfer, the stability of anaerobic microbial communities, and their ability to adapt to environmental change ^4,6,7^. However, there are no accurate methods for measuring the rates at which H_2_ and formate are transferred between microbes or for quantifying interspecies electrical currents in complex communities. Therefore, the importance of HFIT or DIET in microbial communities is typically inferred from the composition of the microbial community.

It has previously been considered that DIET is primarily restricted to environments in which *Geobacter* species are abundant ^8–12^, because *Geobacter* species are the only microbes in pure culture that have been definitively shown to function as electron-donating partners for DIET ^9,13–17^. Studies with *Geobacter metallireducens* have suggested that electrically conductive pili (e-pili) are an important conduit for electron transfer from the electron-donating partner ^9,14,17,18^. It was initially considered that only short pilin monomers (ca. 60 amino acids), such as those found in *G. metallireducens* and *G. sulfurreducens*, could assemble into e-pili ^19^. However, it was subsequently found that larger pilin monomers (> 100 amino aids), phylogenetically distinct from *Geobacter* pilins, can yield e-pili and that e-pili have independently evolved multiple times ^20^. This raises the possibility that some microorganisms not previously known to be capable of extracellular electron transfer may express e-pili to enable DIET.

For example, *Syntrophus aciditrophicus* is one of the most intensively studied pure culture models for HFIT ^21–24^. It was previously concluded that *S. aciditrophicus* would be unlikely to participate in DIET ^24,25^ because it lacks a gene encoding a protein homologous to the Geobacter pilin monomer (PilA) that assembles into electrically conductive pili (e-pili) ^26^.

Here we report that *S. aciditrophicus* expresses e-pili and is capable of growing via DIET. These results, and analysis of the pilin genes of other common syntrophs, indicate that the capacity for DIET should be considered as an option for microorganisms known to grow via HFIT and suggest that DIET may be more widespread than previously considered.

## Results and Discussion

Transmission electron microscopy of *S. aciditrophicus* revealed filaments with a morphology typical of type IVa pili (Fig. 1a). A complement of genes for protein components required for type IV pilus assembly (*pilA, pilD, pilM, pilN, pilO, pilP, pilQ*) and transcriptional control of pilus expression (*pilS, pilR*) are also present in the genome (Supplementary Fig. 1). One gene, SYN 00814 encodes a peptide with a N-terminal domain characteristic of PilA, the pilin monomer for Type IVa pili found in other microrganisms (Fig. 1b). This includes a short leader peptide (13 amino acids) that is cleaved by PilD at the G|FTLIE recognition site and a highly conserved, hydrophobic, pilin assembly interface that facilitates pilin polymerization into pili ^27,28^.

**Fig.1.**
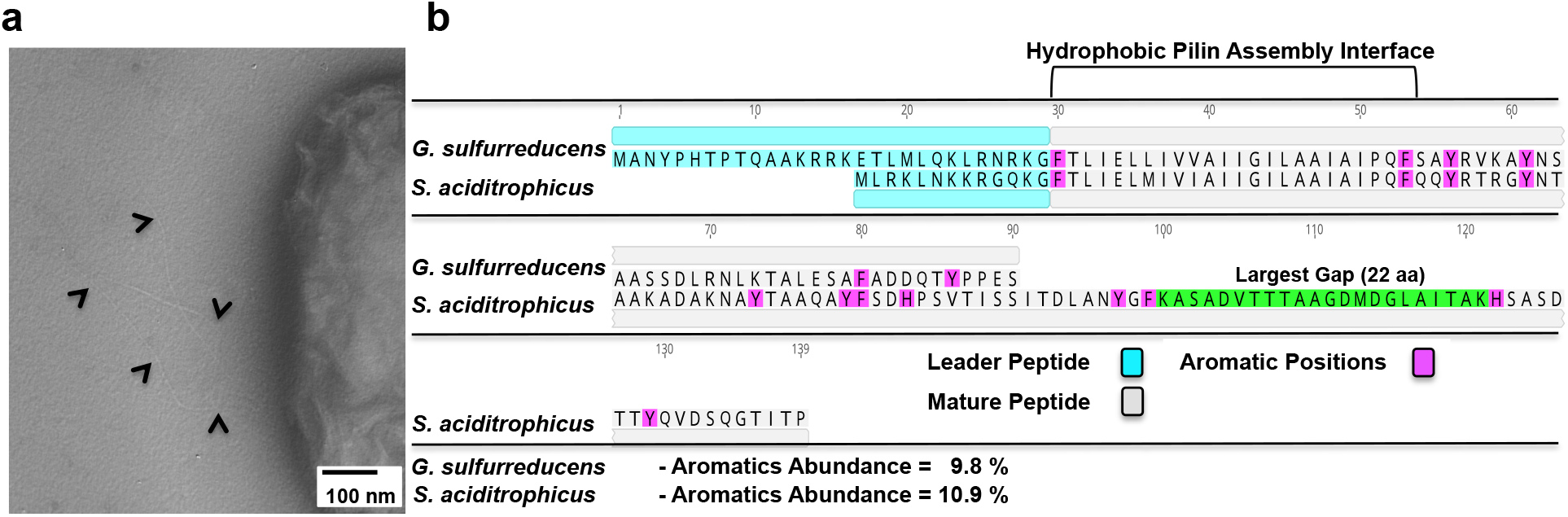
Pili protruding from *S. aciditrophicus* and the putative *S. aciditrophicus* PilA pilin monomer peptide identified from the genome sequence. (**a**) Transmission electron micrograph showing multiple pili protruding from *S. aciditrophicus* (pili highlighted with black arrows). (**b**) Key characteristics of the predicted amino acid sequence of the *S. aciditrophicus* pilin monomer. The PilA monomer of *G. sulfurreducens* that yields conductive pili is shown for comparison. Aromatic amino acid abundance is calculated as a percentage of the mature peptide.

The amino acid sequence of the putative PilA fits the empirical criteria ^20^ for a pilin monomer likely to yield e-pili: 1) aromatic amino acids are located in the key positions required for conductivity in *G. sulfurreducens* e-pili; 2) the abundance of aromatic amino acids (10.9 % of amino acids) is above the minimum threshold of 9 % thought to be necessary for high e-pili conductivity; and 3) no large gaps (> 40 amino acids) that lack aromatic amino acids (Fig. 1b).

Low culture densities of *S. aciditrophicus* prevented harvesting sufficient quantities of pili to measure pili conductance on the electrode arrays previously employed for the study of other e-pili ^20^. Therefore, the method initially employed to document the presence of e-pili in *G. sulfurreducens* ^29^ was adapted ^30^ as an alternative approach. Cultures of *S. aciditrophicus* were directly drop-cast on highly ordered pyrolytic graphite (HOPG), washed with deionized water, and examined with atomic force microscopy. Topographic imaging, (Fig. 2a), indicated that the distance from the surface of the HOPG to the top of the pili (Fig. 2b) was 4.0 + 0.7 nm (n=27; nine different locations on three separate pili). The 4 nm diameter of the *S. aciditrophicus* pili is larger than the 3 nm reported for the e-pili of *Geobacter* species ^31–33^, but thinner than the ca. 5-6 nm diameter of the intensively studied type IVa pili of *Psedumonas* and *Neisseria* species ^34–36^.

**Fig. 2.**
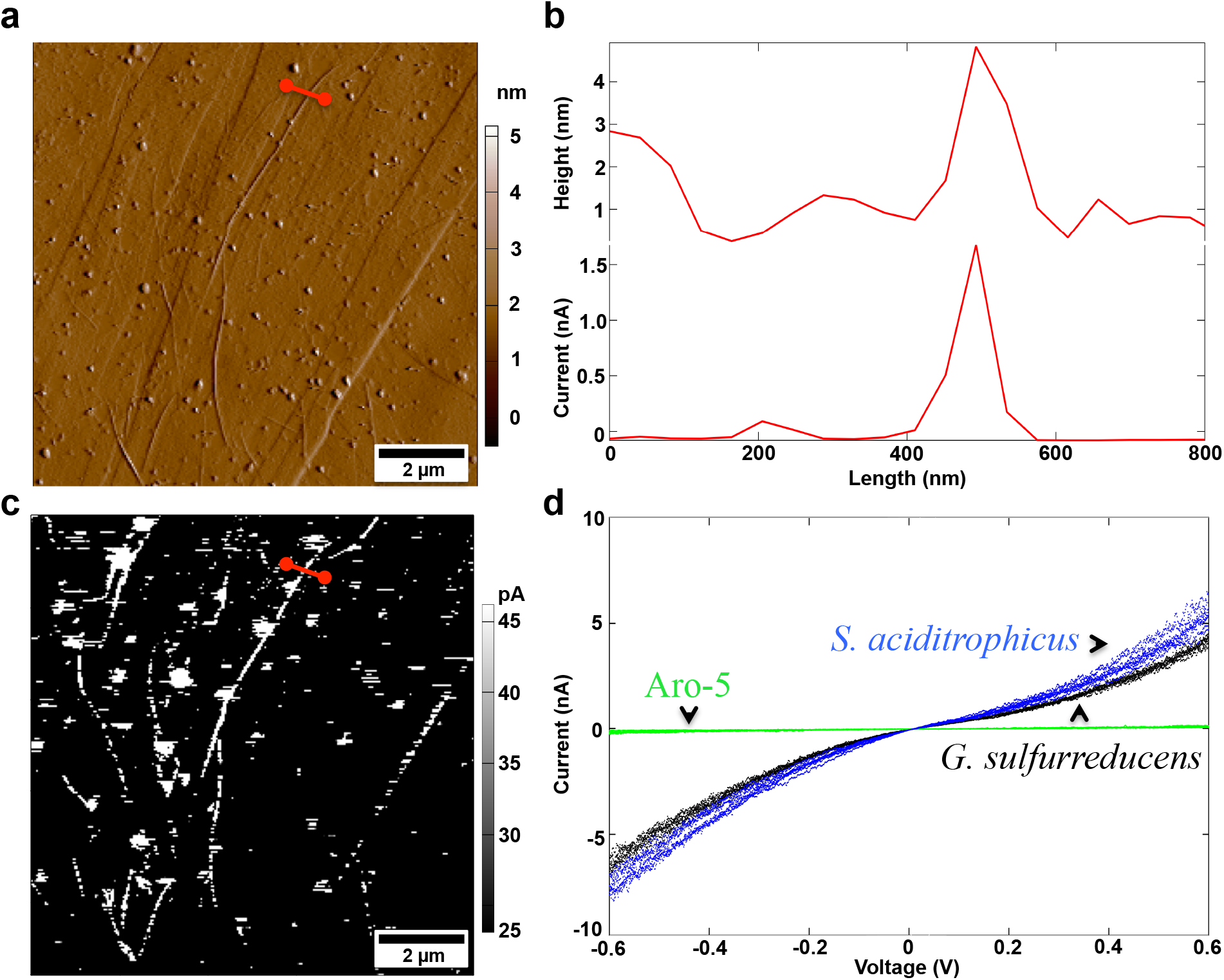
Conducting tip atomic force microscopy demonstrated that *Syntrophus aciditrophicus* pili are electrically conductive. (**a**) Contact mode topographic imaging of pili on highly ordered pyrolytic (HOPG). Red line designates the cross section examined in (b). (**b**) Topographic analysis of the height/diameter of an individual pilus and corresponding current measurements (100 mV differential between the tip and the HOPG) across the pilus cross-section. (**c**) Current response of the pili shown in (a) with an applied 100 mV differential. (**d**) Current-voltage analysis of individual pili of *S. aciditrophicus* (blue data points), wild-type *G. sulfurreducens* (black data points) and the Aro-5 strain of *G. sulfurreducens* (green data points). Current-voltage spectroscopy is shown for one pilus of each type and are representative of analysis of three distinct locations on three separate pili of each type. Additional scans of *S. aciditrophicus* pili available in Supplementary Fig. 2. The *G. sulfurreducens* wild-type and strain Aro-5 data are from reference ^30^.

Conductive imaging revealed that the pili were electrically conductive (Fig. 2b, c). Point-mode current-voltage (I-V) spectroscopy (Fig. 2d; Supplementary Fig. 2) yielded an Ohmic-like response with a conductance of 6.1 ± 1.2 nS (mean ± standard deviation; n=9) that was similar to the previously reported ^30^ conductance of 4.5 ± 0.3 nS for wild-type *G. sulfurreducens* pili. As previously reported ^30^, the pili from *G. sulfurreducens* strain Aro-5, which lack key aromatic amino acids required for high conductivity ^31,33,37,38^, had much lower (0.004 + 0.002 nS) conductance (Fig. 2d).

The lack of tools for genetic manipulation of *S. aciditrophicus* limited further functional analysis of the putative PilA gene predicted to yield its e-pili in the native organism. Therefore, the gene was heterologously expressed in *G. sulfurreducens*, replacing the native *G. sulfurreducens pilA* with an approach that has successfully yielded *G. sulfurreducens* strains that express a diversity of both highly conductive and poorly conductive heterologous pili ^20,32,39–41^. This new *G. sulfurreducens* strain, designated strain SP (for *Syntrophus* pili), expressed abundant pili (Fig. 3a). The heterologously expressed pili had a diameter (4.1 ± 0.4 nm) and conductance (5.9 ± 0.7 nS) nearly identical to the native pili expressed by *S. aciditrophicus* (Fig. 3b). *G. sulfurreducens* strain SP produced electrical current densities comparable to the control strain expressing the *G. sulfurreducens* wild-type *pilA* (Fig. 3c). Such high current densities are only possible when *G. sulfurreducens* expresses e-pili ^20,37^. As previously reported ^20,37^, *G. sulfurreducens* strain Aro-5, with its poorly conductive pili, produced much lower current densities (Fig. 3c). Networks of pili sheared from the electrode-grown biofilm of strain SP, purified, and drop cast on electrode arrays as previously described ^20^, had a conductance comparable to *G. sulfurreducens* wild-type pili networks (Fig. 3d). These results further demonstrated that the *S. aciditrophicus* PilA gene yields a pilin monomer that can assemble into e-pili.

**Fig. 3.**
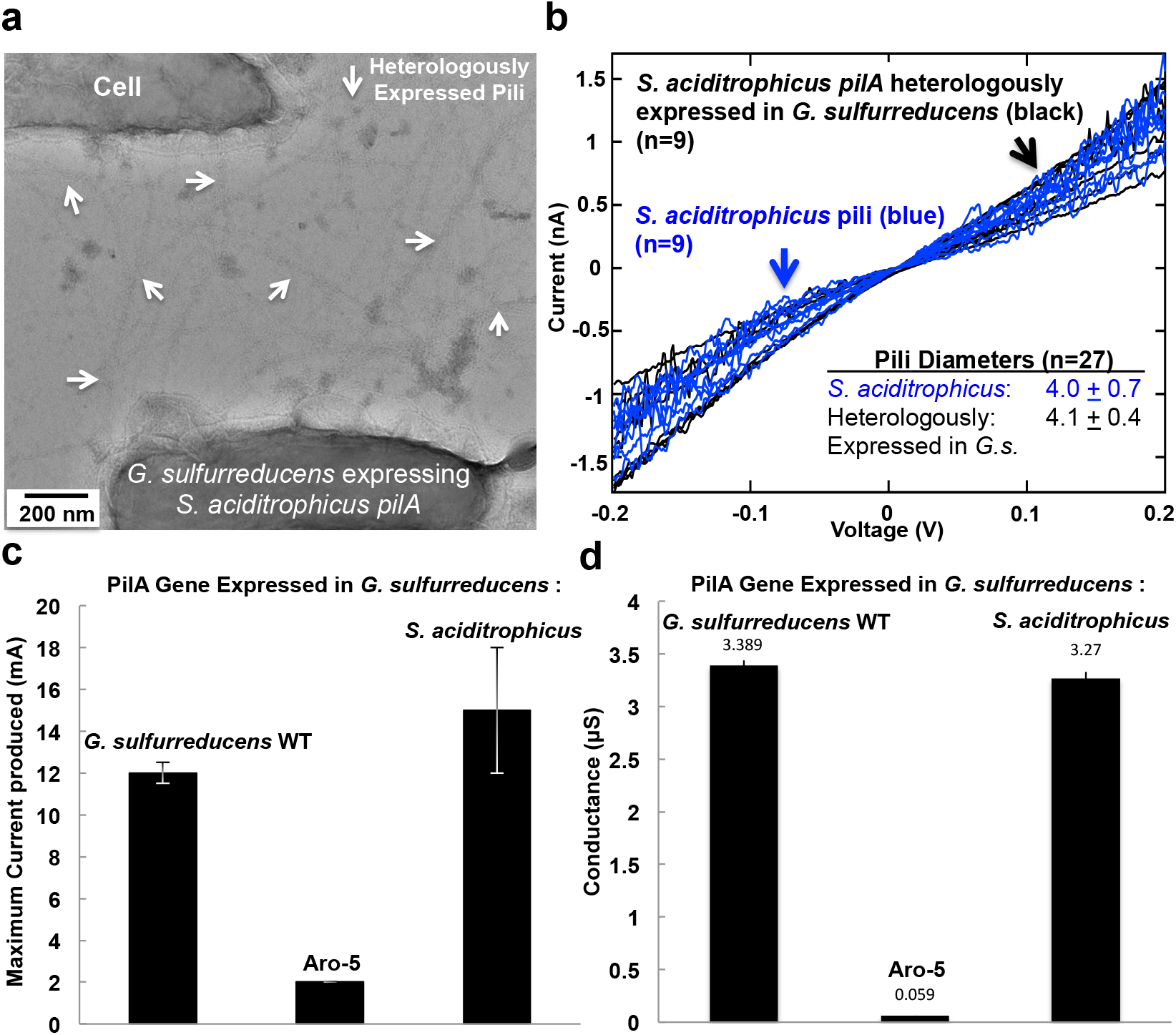
Physical, electrical, and functional, analysis of the putative *Syntrophus aciditrophicus* PilA via heterologous expression in *Geobacter sulfurreducens*. (**a**) Transmission electron micrograph of pili expression in the strain of *G. sulfurreducens* in which the native *pilA* was replaced with the *S. aciditrophicus pilA*. Examples of the location of pili is designated with white arrows. (**b**) Current-voltage profile for three separate locations on three pili expressed by *S. aciditrophicus* pili (data aggregated from individual plots in Fig.1d and Supplementary Fig. S2) and for three separate locations on three pili expressed in the SP strain of *G. sulfurreducens* in which the wild-type *pilA* was replaced with the *S. aciditrophicus pilA* (individual plots shown in Supplentary Fig. 3). (**c**) Current production by *G. sulfurreducens* with wild-type PilA; the synthetic Aro-5 PilA designed to yield poorly conductive pili; or *S. aciditrophicus* PilA. (**d**) Four-probe conductance (mean ± standard deviation n=9) of films of purified pili from strains of *G. sulfurreducens* expressing; *S. aciditrophicus* PilA compared with previously reported ^20^ conductances of pili derived from wild-type and Aro-5 strains of *G. sulfurreducens.*

The presence of e-pili in *S. aciditrophicus* suggested it might be capable of establishing an electrical connection for DIET. To evaluate this, *S. aciditrophicus* was grown in co-culture with *G. sulfurreducens*, the microbe most intensively studied as an electron-accepting partner for DIET ^4^. *G. sulfurreducens* can also function as a H_2_- and formate-consuming partner for HFIT ^42^. Therefore, even if *S. aciditrophicus* was unable to grow via DIET it would be expected to grow in co-culture with *G. sulfurreducens* via HFIT, providing a positive control to ensure that the culture medium was appropriate for co-culture growth. The electron donor was benzoate, a substrate that *S. aciditrophicus* can metabolize, but *G. sulfurreducens* cannot. The electron acceptor was fumarate, an electron acceptor that only *G. sulfurreducens* can utilize. In the presence of a H_2_/formate-consuming partner, *S. aciditrophicus* metabolizes benzoate to acetate with the production of either H_2_:

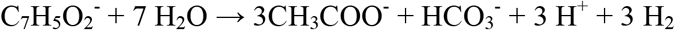

or formate:

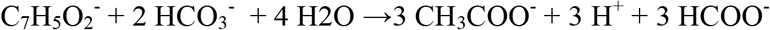

With electron transfer via DIET the relevant reaction is:

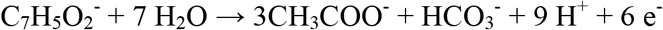

In addition to oxidizing H_2_ and formate, or consuming electrons released during DIET, *G. sulfurreducens* can also metabolize acetate, making the overall reaction expected for the oxidation of benzoate with the reduction of fumarate to succinate in the co-culture:

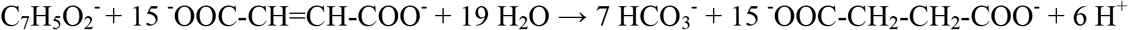

*S. aciditrophicus/G. sulfurreducens* co-cultures grew with repeated sub-culturing and, within the experimental error, exhibited the expected stoichiometry of benzoate consumption and succinate production (Fig. 4a). In these co-cultures HFIT, DIET, or a combination of the two, were feasible. Therefore, to eliminate the possibility of HFIT, *S. aciditrophicus* was co-cultured with the previously described strain of *G. sulfurreducens* ^18^, designated here as *G. sulfurreducens*_HF_, that cannot utilize H_2_ or formate because the genes for the uptake hydrogenase and formate dehydrogenase were deleted. The *S. aciditrophicus/ G. sulfurreducens*_HF_ co-culture had a longer initial lag period, but then metabolized benzoate with the reduction of fumarate almost as fast as the co-culture with wild-type *G. sulfurreducens* (Fig. 4b). These results demonstrate that *S. aciditrophicus* can grow via DIET.

**Fig. 4.**
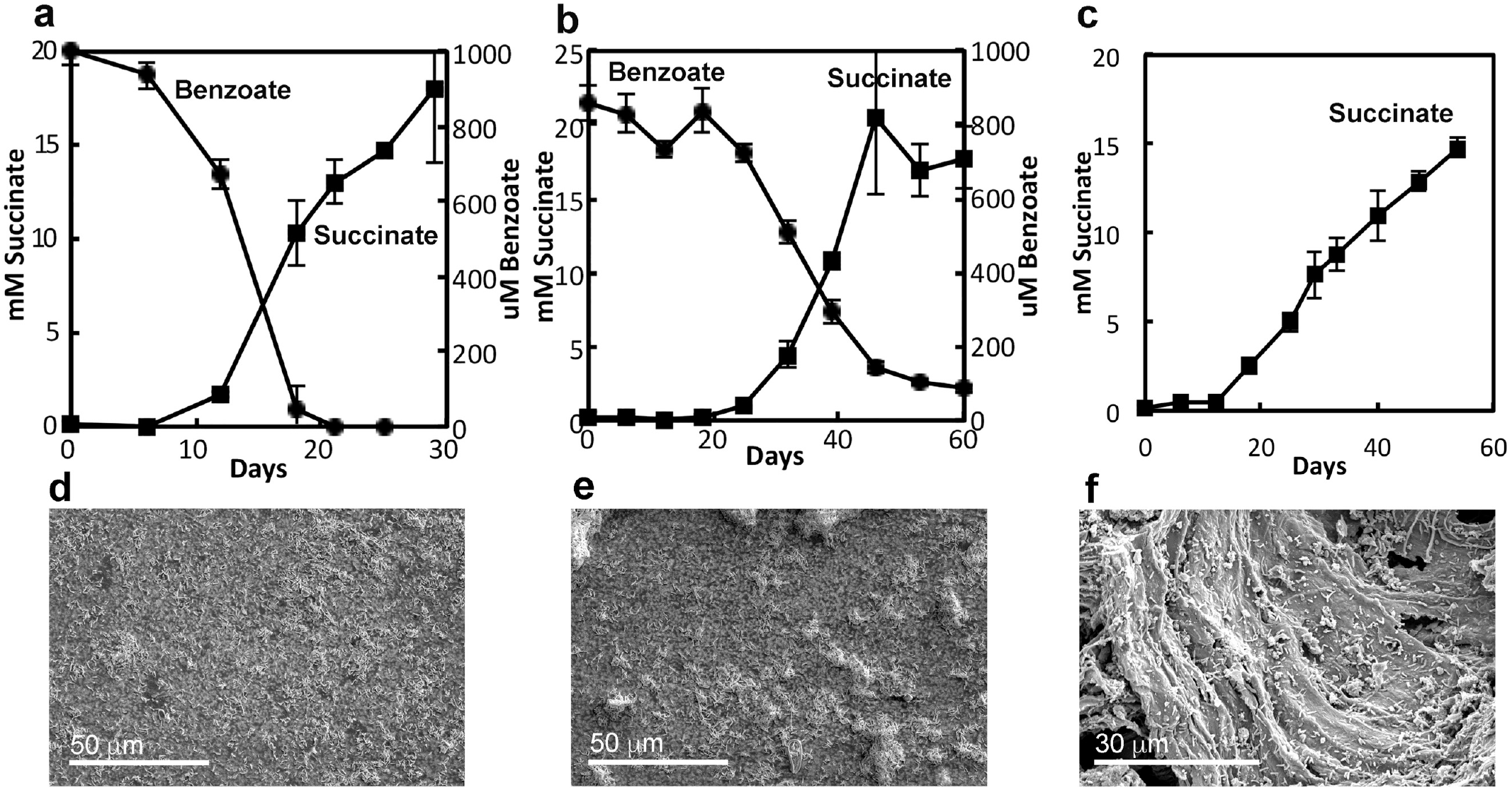
Co-cultures of *Syntrophus aciditrophicus* and *Geobacter sulfurreducens* grow via DIET. Metabolism with *S. aciditrophicus* co-cultured with (**a**) wild-type *G. sulfurreducens*, (**b**) *G. sulfurreducens*_HF_, which is unable to use H_2_ or formate. (**c**) *G. sulfurreducens*_HF_ with granular activated carbon (GAC) amendment. No acetate or formate was detected in any of the co-cultures. GAC interfered with determination of benzoate, which is not shown for GAC-amended cultures. Scanning electron micrographs (SEM) of cells collected on filters from co-cultures with (**d**) *G. sulfurreducens* wild-type or (**e**) *G. sulfurreducens* strain_HF_ demonstrating greater aggregation in co-cultures with *G. sulfurreducens*_HF_ in which DIET was the only option for interspecies electron exchange. (**f**) SEM of cells on GAC from co-culture of *S. aciditrophicus* with *G. sulfurreducens* strain_HF_. Circles-benzoate; squares-succinate. Data are the mean and standard deviation of triplicate cultures.

DIET requires physical electrical contacts between the electron-donating and electron-accepting partner, whereas contact is not required for HFIT ^4,5^. In some co-cultures the requirement for contact manifests as visible aggregates ^13^, but other DIET co-cultures produce small, relatively fragile aggregates ^14^. There were no visible aggregates in the *S. aciditrophicus/G. sulfurreducens* co-cultures. However, more small clumps were seen in scanning electron micrographs of co-cultures in which *G. sulfurreducens*_HF_ was the electron-accepting partner than co-cultures with wild-type *G. sulfurreducens* provided as the electron-accepting partner (Fig. 4 d,e). This observation is consistent with the need for contact between electron-donating and electron-accepting partners participating in DIET. In contrast, HFIT was an option for the co-cultures established with wild-type *G. sulfurreducens* and aggregation is not required when *G. sulfurreducens* grows via HFIT ^42^.

Granular activated carbon (GAC) can greatly reduce the initial lag time in establishing DIET-based co-cultures because both partners attach to GAC, which functions as an electrically conductive conduit ^14,43^. GAC substantially reduced the lag time of the *S. aciditrophicus/ G. sulfurreducens*_HF_ co-cultures (Fig. 4c). As previously observed for other co-cultures in which GAC promoted DIET, cells from the *S. aciditrophicus/ G. sulfurreducens*_HF_ co-culture heavily colonized the GAC (Fig. 4e), consistent with a GAC conduit for DIET.

In addition to e-pili, multi-heme outer-surface *c*-type cytochromes appear to play an important role in DIET in *Geobacter* species ^4,5^. However, the S. *aciditrophicus* genome encodes only a few putative *c*-type cytochromes ^24^ and cytochromes were not readily apparent in heme-stained preparations of cell protein (Supplementary Fig. 4). Not all microbes require cytochromes for effective electron transport to the outer cell surface ^44^. More detailed examination of the role of e-pili and other *S. aciditrophicus* components in DIET will require the development of methods for genetic manipulation of this microorganism.

*S. aciditrophicus* is the first isolate outside the genus *Geobacter* found to grow via DIET and it is the first syntroph shown to have the option to grow via HFIT or DIET. Other *Syntrophus* species also have pilin genes with the empirical criteria, detailed above, of aromatic amino acid placement and abundance likely to yield e-pili (Fig. 5). The pilins of other diverse genera of syntrophic microorganisms known to grow via HFIT in defined co-cultures also meet these criteria (Fig. 5). Establishing conditions that favor DIET often enrich for microbes in these genera ^6^. For example, in enrichment cultures specifically designed to promote the metabolism of propionate or butyrate via DIET, *Smithella* (propionate enrichment) or *Syntrophomonas* (butyrate enrichment) species were the most abundant bacteria ^45–47^. The greater energetic demands ^48^ required for synthesizing the abundant aromatic amino acids that are required for e-pili conductivity suggests that e-pili provide a strong selective advantage under some environmental conditions. Conferring the capacity for DIET is a likely explanation.

**Fig. 5.**
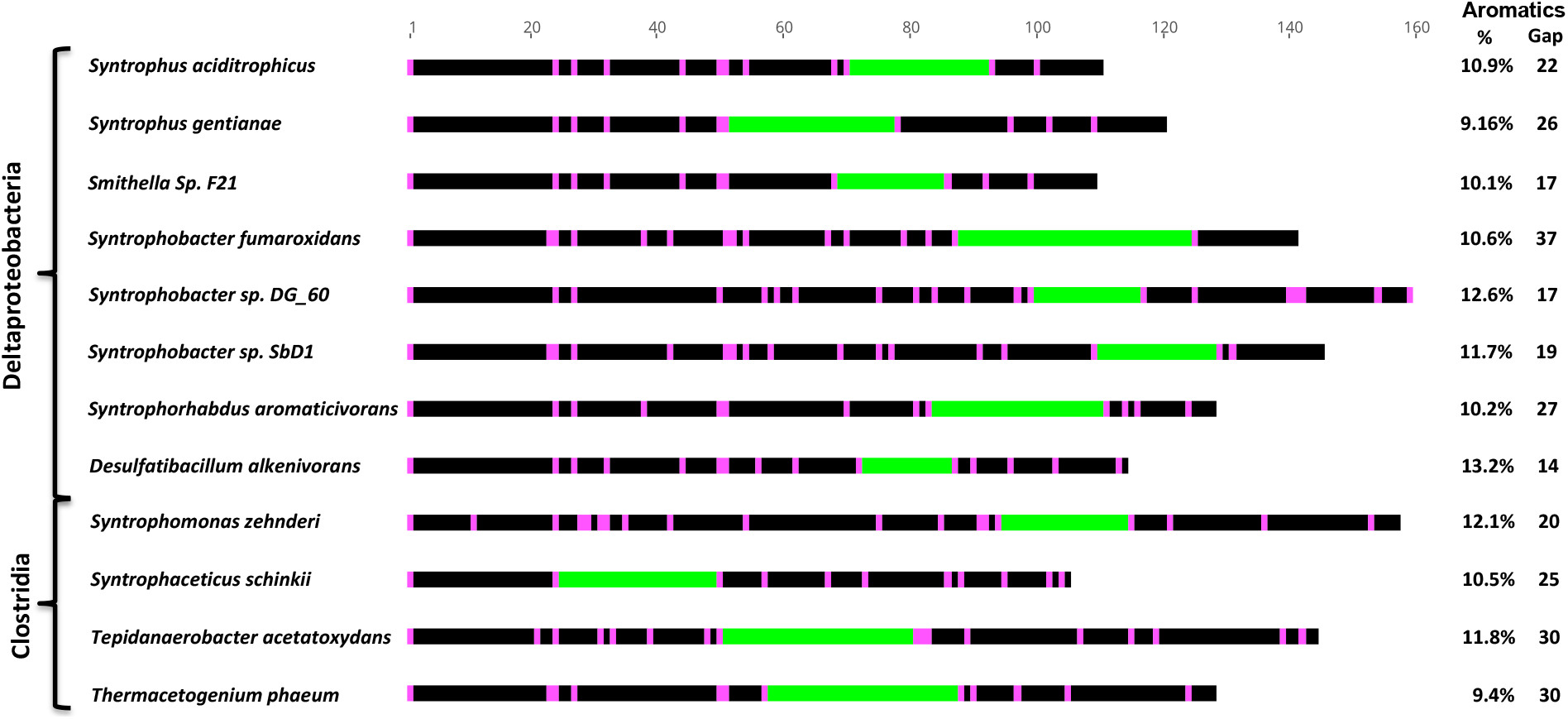
A diversity of syntrophs known to produce H_2_ and/or formate as an interspecies electron carrier encode for pilin monomers likely to yield electrically conductive pili. Position of aromatic amino acids designated in magenta and the largest gap that lacks aromatic amino acids designated in green. The full protein sequence alignment for each pilin can be found in Supplementary Fig. 5. Each of the pilins meets the empirical criteria derived from previous studies ^20^ of: 1) aromatic amino acids located in the key positions required for conductivity in *G. sulfurreducens* e-pili; 2) the abundance of aromatic amino acids greater than 9%; and 3) no aromatic-free gaps of greater than 40 amino acids.

For decades, interactions within complex anaerobic microbial communities have been interpreted through the lens of HFIT, but the available data does not rule out the possibility of DIET, which was not known to be an option at the time. For example, estimated H_2_ turnover rates in anaerobic digesters, rice paddy soils, and aquatic sediments were consistently less than 10 % of the independently determined rate of methane production derived from the reduction of carbon dioxide to methane ^49–51^, a result that is consistent with DIET providing most of the electrons for carbon dioxide reduction. The mismatch between measured H_2_ turnover rates and the concept of H_2_ as a primary interspecies electron carrier was rationalized with the suggestion that there was a separate pool of H_2_ within closely juxatapositioned assemblages of H_2_ producers and H_2_ consumers. However, the existence of two distinct pools of H_2_ that did not equilibrate over time via diffusion, a seemingly physical impossibility, was never verified. Attempts to accurately measure formate fluxes have also been problematic ^52^ and there is no method for directly measuring DIET-based electron fluxes in complex communities.

A fresh perspective and new analytical tools will be required to resolve the relative importance of HFIT and DIET in diverse anaerobic microbial communities. Just as electron-accepting partners have different gene expression patterns depending on whether they are participating in HFIT or DIET ^18,53^, it may be possible to determine whether electron-donating syntrophs are engaged in DIET or HFIT from metatranscriptomic analysis of anaerobic communities.

## METHODS

### Bacterial strains, plasmids and culture conditions

*Syntrophus aciditrophicus, G. sulfurreducens* wild-type, *G. sulfurreducens* strain Aro-5, and *G. sulfurreducens*_HF_ were obtained from our laboratory culture collections. A strain of *G. sulfurreducens* expressing *pilA* from *S. aciditrophicus* rather than native *G. sulfurreducens pilA* was constructed as previously described ^20^. *S. aciditrophicus* and *G. sulfurreducens* strains were routinely grown under strict anaerobic conditions at 30 °C in previously described ^54^ defined, bicarbonate-buffered medium with N_2_:CO_2_ (80:20) as the gas phase. For *S. aciditrophicus*, the medium was amended with crotonate (20 mM) and for *G. sulfurreducens* strains, acetate (10 mM) was the electron donor and fumarate (40 mM) was the electron acceptor. The presence of *c*-type cytochromes in whole cell lysates was evaluated with heme-staining of proteins separated on denaturing polyacrylamide gels as previously described ^17^. *G. sulfurreducens* strains were grown with a graphite electrode as the electron acceptor as previously described ^55^.

Cocultures were established in 10 ml of culture medium ^54^ in anaerobic pressure tubes with benzoate (1 mM) as the electron donor and fumarate (40 mM) as the electron acceptor with cysteine (2 mM) and sulfide (1 mM) added as reducing agents. When noted, cultures were amended with granular activated carbon (0.25 g; 8-20 mesh). Benzoate, acetate, and succinate were analyzed with high-performance liquid chromatography as previously described ^16^. Previously described methods were employed for transmission electron microscopy ^21^ and scanning electron microscopy ^26^.

### Pili conductivity

A 100 μl sample of cultures was drop cast onto highly oriented pyrolytic graphite (HOPG). After 10 min, the HOPG was washed twice with 100 μl of deionized water, blotted dry to remove excess water, and dried for 12 hours at 24 °C in a desiccator. Samples were equilibrated with atmospheric humidity for at least 2 hours and then examined with an Oxford Instruments Cypher ES Environmental AFM in ORCA electrical mode equipped with a Pt/Ir-coated Arrow-ContPT tip with a 0.2 N/m force constant (NanoWorld AG, Neuchâtel, Switzerland). Pili were located in contact mode, with a set point of 0.002 V and a scan rate of 1.5 Hz. For conductive imaging, the grounded tip, attached to a transimpedance amplifier, served as a translatable top electrode to locally detect the current response of the individual pili to a 100 mV bias applied to the HOPG substrate. Individual pili conductivity was further characterized by lightly pressing the AFM tip (set point 0.002 V) to the top of the pili and applying a quadruplicate amplitude of ±0.6 V voltage sweep at a frequency of 0.99 Hz, receiving ca. 8,000 points of reference per measurement. Pili expressed in *G. sulfurreducens* were further analyzed with four probe conductivity measurements on films of pili purified from cells as previously described ^20^.

