## Supplementary Files for "*Syntrophus* Conductive Pili Demonstrate that Common Hydrogen-Donating Syntrophs can have a Direct Electron Transfer Option"

**Supplementary Figures.**

David J.F. Walker[1,2], Kelly P. Nevin[1], Dawn E. Holmes[1,3], Amelia-Elena Rotaru[1,4], Joy E. Ward[1], Trevor L. Woodard[1], Jiaxin Zhu[5], Toshiyuki Ueki[1], Stephen S. Nonnenmann[2,5], Michael J. McInerney[6], and Derek R. Lovley[1,2*]

[1]Department of Microbiology, University of Massachusetts-Amherst, Amherst, MA, USA
[2]Institute for Applied Life Sciences, University of Massachusetts-Amherst
[3]Department of Physical and Biological Science, Western New England University, Springfield, MA, USA
[4]Department of Biology, University of Southern Denmark, Odense, Denmark
[5]Department of Mechanical and Industrial Engineering, University of Massachusetts-Amherst
[6]Department of Microbiology and Plant Biology, University of Oklahoma, Norman, OK, USA

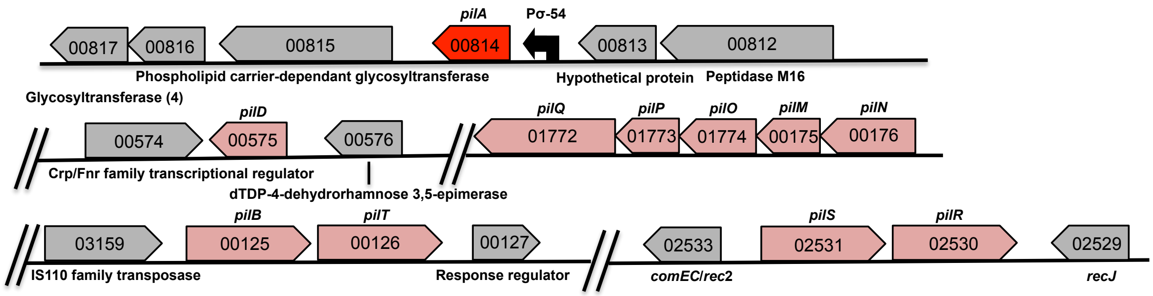


**Supplementary Figure 1. Location of the pilin assembly genes within the chromosome of _S. aciditrophicus_.** The putative PilA gene, shown in Dark Red, is downstream of a Sigma 54 promoter likely to be controlled by the PilR/PilS two-component system. The other pilin assembly genes are shown in light red.

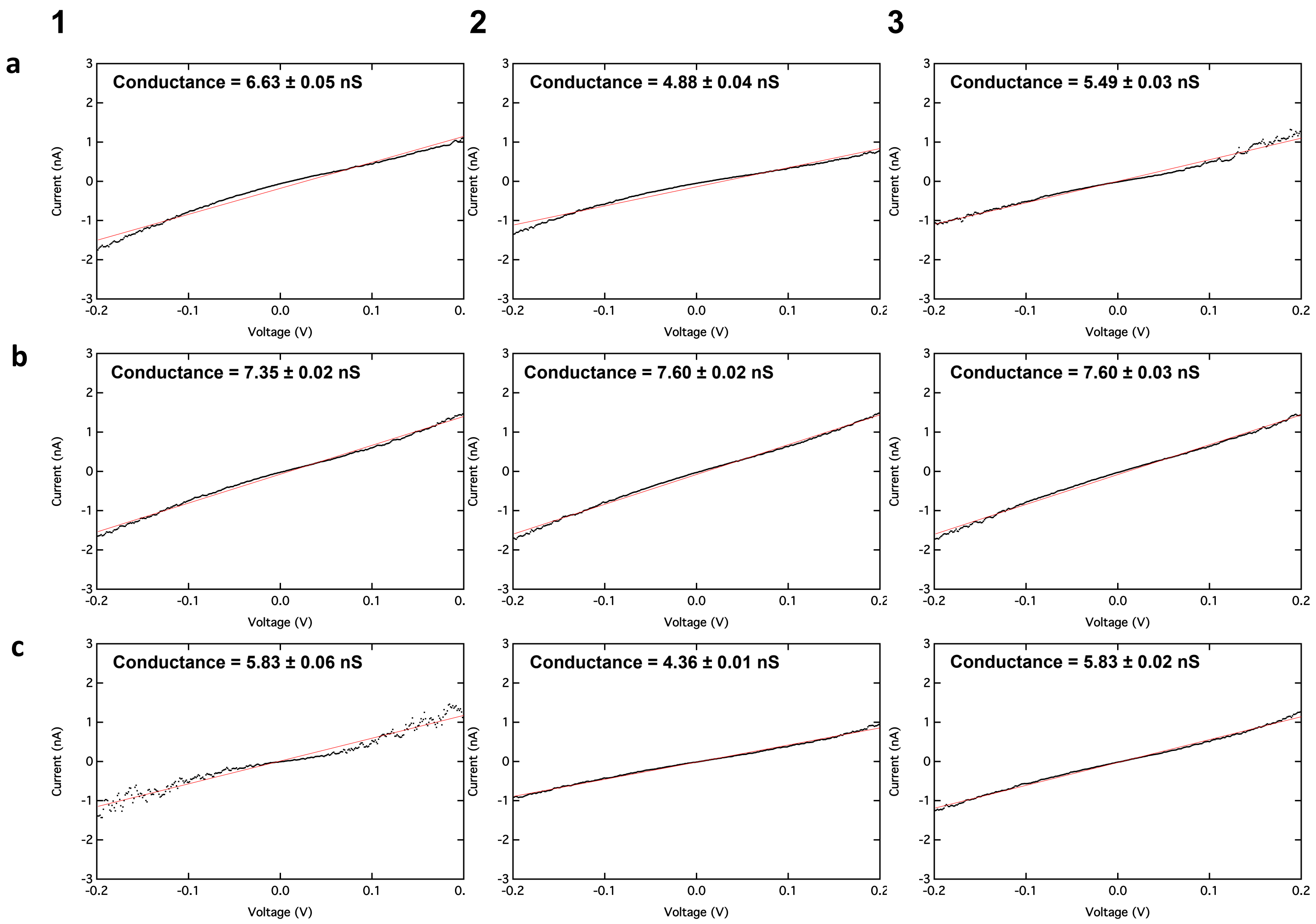


**Supplementary Figure 2. Point-mode current response (I-V) spectroscopy measurements and conductance calculations of the three *S. aciditrophicus* pili, shown in Fig 1d and Fig 3b of the primary text. (a)** Shown in Panel a1 are the measurements of the pili analyzed in Figure 1d of the primary text. Panel a2 and a3 are two more independent measurements at different locations on the same pilus. **(b & c)** Measurements on two additional pili at three different locations for each pilus. Conductance calculated from a linear fit model between -0.2 V and 0.2 V. Units are nano-Siemens (nS)

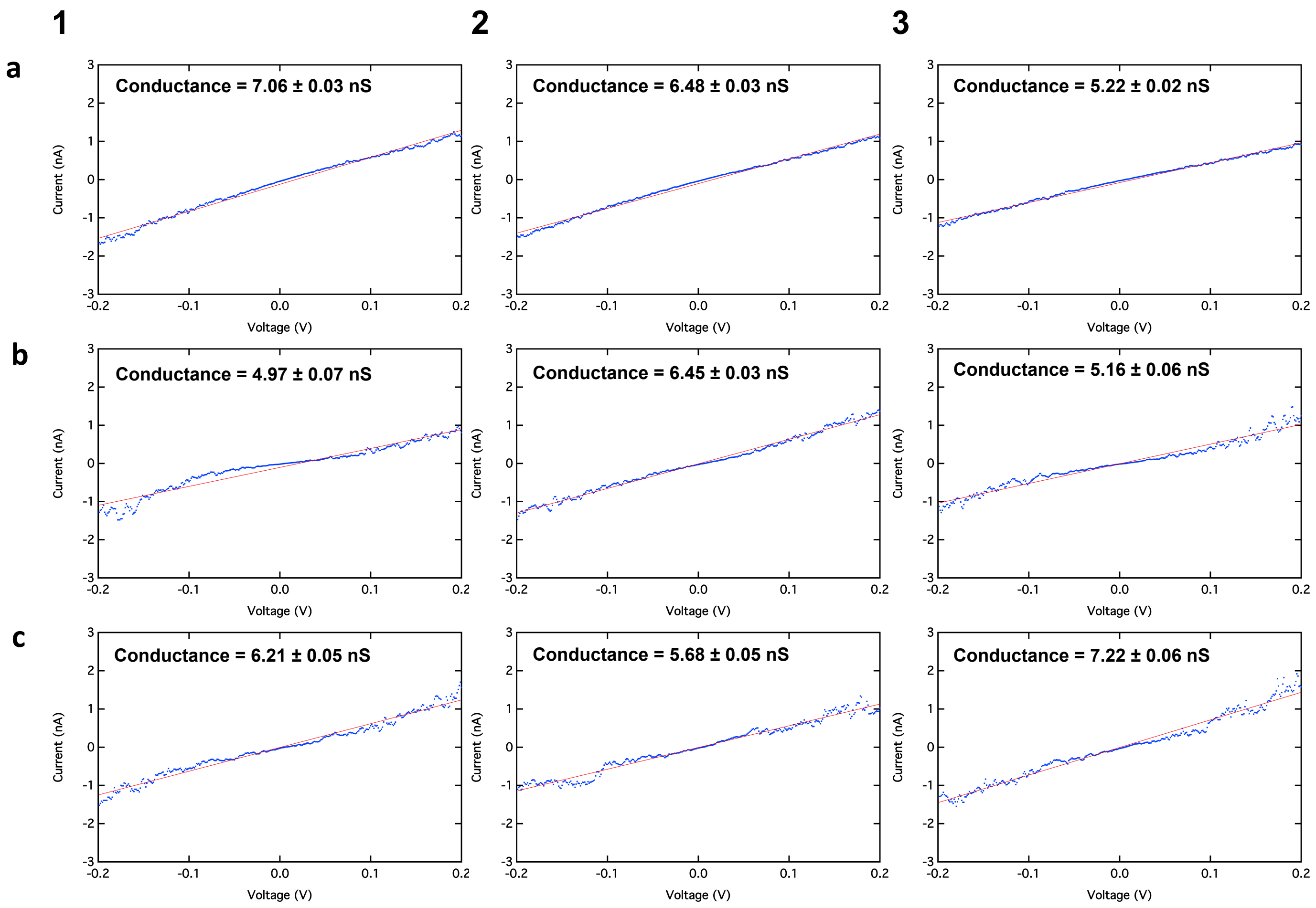


**Supplementary Figure 3. Point-mode current response (I-V) spectroscopy measurements and conductance calculations for the three pili from *G. sulfurreducens* strain SP, shown in Fig 3b of the primary text.** Letters designate the individual pili, numbers designate the individual points on that pili. Calculations were made using a linear fit model between -0.2 V and 0.2 V. Units are nano-Siemens (nS)

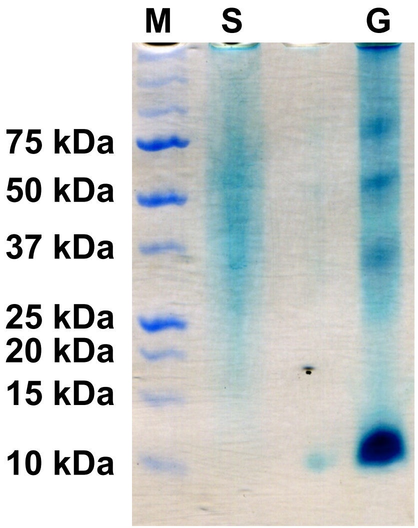


**Supplementary Figure 4. Heme-staining of SDS PAGE gels of whole cell extracts of *Syntrophus aciditrophicus* (S) and *Geobacter sulfurreducens* (G).** Despite adding 120-fold more *S. aciditrophicus* cell protein (12 µg) than *G. sulfurreducens* cell protein (0.1µg), *c*-type cytochromes were not readily detected in *S. aciditrophicus*. M designates a lane for protein standards of the sizes designated on the left.

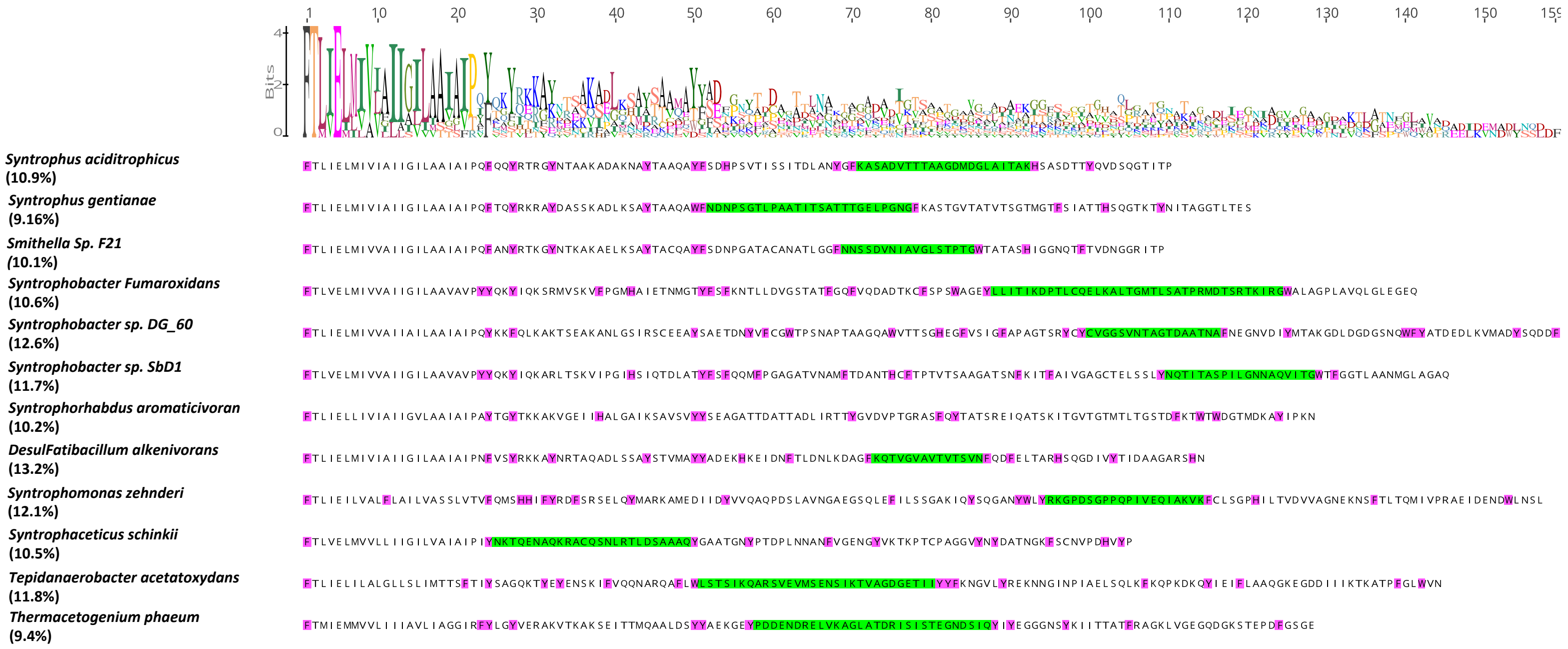


**Supplementary Figure 5. Amino acid alignment of the pilins of the syntrophs in Fig. 5 of the primary text.** Highlighted in pink are the positions of the aromatic amino acids. Highlighted in green are the 2 largest gaps. At the top is the logo sequence alignment showing the high conservation at the N-terminal and at specific aromatic amino acids positions in the first 55 amino acids of the mature peptide. Percentage of aromatics in the mature peptide is listed below the organism's name.
